# Antiviral activity of bacterial TIR domains via signaling molecules that trigger cell death

**DOI:** 10.1101/2021.01.06.425286

**Authors:** Gal Ofir, Ehud Herbst, Maya Baroz, Daniel Cohen, Adi Millman, Shany Doron, Nitzan Tal, Daniel B. A. Malheiro, Sergey Malitsky, Gil Amitai, Rotem Sorek

## Abstract

The Toll/interleukin-1 receptor (TIR) domain is a canonical component of animal and plant immune systems. In plants, intracellular pathogen sensing by immune receptors triggers their TIR domains to generate a molecule which is a variant of cyclic ADP-ribose (v-cADPR). This molecule is hypothesized to activate plant cell death via a yet unresolved pathway. TIR domains were recently also shown to be involved in a bacterial anti-phage defense system called Thoeris, but the mechanism of Thoeris defense remained unknown. In this study we report that phage infection triggers Thoeris TIR-domain proteins to produce an isomer of cyclic ADP-ribose. This molecular signal activates a second protein, ThsA, which then depletes the cell of the essential molecule nicotinamide adenine dinucleotide (NAD) and leads to abortive infection and cell death. We further show that similar to eukaryotic innate immune systems, bacterial TIR-domain proteins determine the immunological specificity to the invading pathogen. Our results describe a new antiviral signaling pathway in bacteria, and suggest that generation of intracellular signaling molecules is an ancient immunological function of TIR domains conserved in both plant and bacterial immunity.

## Introduction

In both animal and plant immune systems, Toll/interleukin-1 receptor (TIR) domains serve as the signal-transducing components of immune receptors that recognize molecular elements of invading pathogens. In humans, TIR domains in Toll-like receptors transfer the signal through protein-protein interactions^1^. In plants, multiple types of immune receptors contain TIR domains, including TIR nucleotide-binding leucine rich repeat proteins (TIR-NLR) that are abundantly present in multiple copies in plant genomes^2^. Recently, plant TIR-containing immune receptors were shown to possess an enzymatic activity, and were reported to produce a variant of cyclic adenine diphosphate ribose molecule (v-cADPR) upon pathogen recognition^3,4^. Activation of plant TIRs leads to a form of cell suicide known as the hypersensitive response, which prevents pathogen propagation^5^. The mechanism through which v-cADPR production promotes cell death in plants is currently unknown^6,7^.

TIR domains were also recently found to serve as essential components in a common prokaryotic immune system called Thoeris, which defends bacteria against phage infection^8^. Thoeris comprises two core proteins, one of which (named ThsB) has a TIR domain (Figure 1A). Although Thoeris was found in thousands of sequenced bacterial and archaeal genomes, its mechanism of action remained unresolved. In the current study we set out to understand the mechanism of Thoeris-mediated immunity against phages, and to trace possible functional and evolutionary connections between bacterial and eukaryotic TIRs.

**Figure 1.**
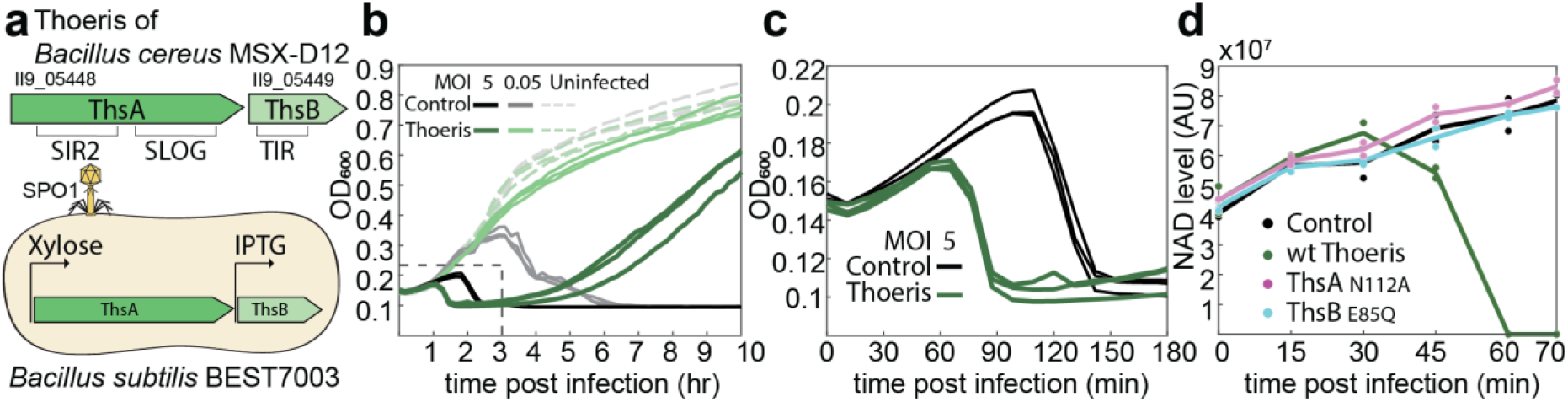
Thoeris causes abortive infection and NAD depletion. **(a)** The Thoeris system of *B. cereus* MSX-D12 used in this study. Gene locus tags and protein domains are indicated (top). Thoeris genes were cloned by genomic integration into *B. subtilis* BEST7003 under the control of inducible promoters (bottom). ThsA was induced with 0.2% xylose and ThsB with 5 μM IPTG. **(b)** Growth curves of Thoeris-expressing (green) and control cultures (black) with and without infection by phage SPO1 at multiplicity of infection (MOI) of 5 or 0.05. Curves represent three independent experiments. **(c)** Magnification of the area marked by a dashed rectangle in panel **b**, representing the first 3 hours of infection by phage SPO1 at an MOI of 5. **(d)** NAD levels in control culture (black), cells expressing wild type Thoeris (green), or mutated Thoeris systems (ThsA_N112A_ mutation in magenta and ThsB_E85Q_ in cyan) during infection. Time 0 represents uninfected cells. Each line represents the mean of two independent experiments, with individual data points shown. Cells were infected by phage SPO1 at an MOI of 5. NAD levels were measured by LC-MS and calculated from the area under the curve of the identified NAD peak at m/z 664.1182 (positive ionization mode) and retention time of 7.8 min. The NAD Peak was identified using a synthetic standard run.

## Results

### Thoeris leads to abortive infection via cellular NAD depletion

We first asked whether Thoeris defense leads to cell death akin to plant TIR-containing immune systems. Multiple bacterial defense systems are known to protect bacteria by forcing them to commit suicide once infected by phage, a process known as abortive infection^9^. Abortive infection systems lead to death of the infected cell prior to the production of new phage particles, thus preventing phage replication and spread within the bacterial culture. To test if Thoeris is an abortive infection system, we carried out experiments using *Bacillus subtilis* cells expressing the Thoeris system of *Bacillus cereus* MSX-D12^8^ (Figure 1A). Cells expressing Thoeris proteins from inducible promoters were protected against phage SPO1 in plaque assay experiments (Methods; Figure 1A, Figure S1A-B). Thoeris-mediated protection from SPO1 infection was also observed in liquid cultures, if phages were added to the culture in low multiplicity of infection (MOI) (Figure 1B). However, adding phages in high multiplicity of infection, in which nearly all cells are expected to be infected by the initial phage inoculum, resulted in a premature culture collapse (Figure 1B-C) that did not involve release of new phages (Figure S1C). The collapse of the culture occurred 70–80 minutes following initial infection, which is earlier than phage-induced lysis that was observed in Thoeris-lacking cells after 120 minutes (Figure 1C). These results suggest that Thoeris defends via abortive infection, leading to death of cells infected by phage SPO1 prior to the maturation of phage progeny.

Although Thoeris-expressing cells demonstrated premature collapse as compared to control cells, the Thoeris culture was able to recover following the collapse, in contrast to control cells that did not recover following phage lysis (Figure 1B). This may be due to a subpopulation of Thoeris-expressing cells that were not infected and therefore able to grow once phages were eliminated during infection of the other cells, or a subpopulation that did not completely lyse following Thoeris activation.

The Thoeris defense system is comprised of two genes, *thsA* and *thsB*, and the presence of both of them is essential for Thoeris defense^8^. ThsB contains a TIR domain, and ThsA most commonly contains a sirtuin (SIR2) domain at its N-terminus^10^ (Figure 1A). It was recently shown that the SIR2 domain within ThsA has a catalytic NADase activity^11^. As NAD depletion has been recently implicated in abortive infection in other phage defense systems^12^, we hypothesized that NAD depletion by ThsA is involved in Thoeris abortive infection. To test this, we monitored NAD levels during phage challenge using liquid chromatography and mass spectrometry (LC-MS). In Thoeris-expressing cells, a complete depletion of NAD was observed 60 minutes after the onset of infection by phage SPO1 (Figure 1D), temporally preceding the culture collapse that occurred 70–80 minutes post infection (Figure 1C). The product of NAD cleavage, ADP-ribose, was detected simultaneously with the decline of NAD levels (Figure S1D). Such NAD depletion and ADP-ribose production were not observed in control cells lacking Thoeris, or in uninfected Thoeris-containing cells (time 0) (Figures 1D, S1D). A point mutation in the active site of the NADase domain in ThsA (N112A), which was shown to be essential for its activity *in vitro*^11^, abolished Thoeris-mediated defense^8^ (Figure S1). The same mutation also abrogated NAD depletion following infection (Figure 1D), suggesting that the NADase activity of ThsA is responsible for the observed NAD depletion.

### Thoeris TIR proteins produce a signaling molecule that activates the ThsA NADase

The TIR domain in the ThsB protein was previously shown to be essential for Thoeris defense^8^, and, accordingly, a mutation in the conserved glutamic acid of the ThsB TIR domain (E85Q) abolished phage defense and NAD depletion (Figures 1D, S1). A mutation in the parallel conserved glutamic acid in plant TIR domain proteins prevented the production of the v-cADPR molecule following recognition of pathogen molecular signatures^3,4^. We therefore hypothesized that the Thoeris TIR protein may generate, in response to phage infection, a signaling molecule akin to the v-cADPR molecules generated by plant TIRs. Under this hypothesis, the signaling molecule generated by the Thoeris TIR-domain protein may trigger the NADase activity of ThsA, which would then lead to abortive infection.

To test this hypothesis, we expressed only the Thoeris ThsB TIR protein in *B. subtilis* cells, and subjected these cells to infection by phage SPO1. At several time points during infection, we lysed the infected cells and filtered the lysates to include only molecules smaller than 3 kDa. We incubated purified ThsA protein with these lysates *in vitro* to test if the lysates affect the NADase activity of ThsA (Figure 2A). To this end, we used a reporter for NAD cleavage, nicotinamide 1,N6-ethenoadenine dinucleotide (εNAD), which generates a fluorescent signal following cleavage by NADases^13^. Purified ThsA showed marked NADase activity when incubated with cell lysates derived from infected cells that expressed the TIR-containing ThsB protein (Figure 2B). Only lysates derived from cells 60 or 70 minutes after infection triggered the NADase activity of ThsA (Figure 2B, Figure S2A), temporally coinciding with the NAD depletion observed in infected cells expressing Thoeris *in vivo* (Figure 1D). Under the experimental conditions used, ThsA showed no NADase activity when incubated with cell lysates from uninfected cells, or with lysates from infected control cells that did not express the ThsB TIR protein, or with lysates from infected control cells that expressed the ThsBE85Q mutant (Figures 2B, S2A). These results suggest that during phage infection, Thoeris TIR proteins generate a small molecule capable of activating the NADase activity of ThsA.

**Figure 2.**
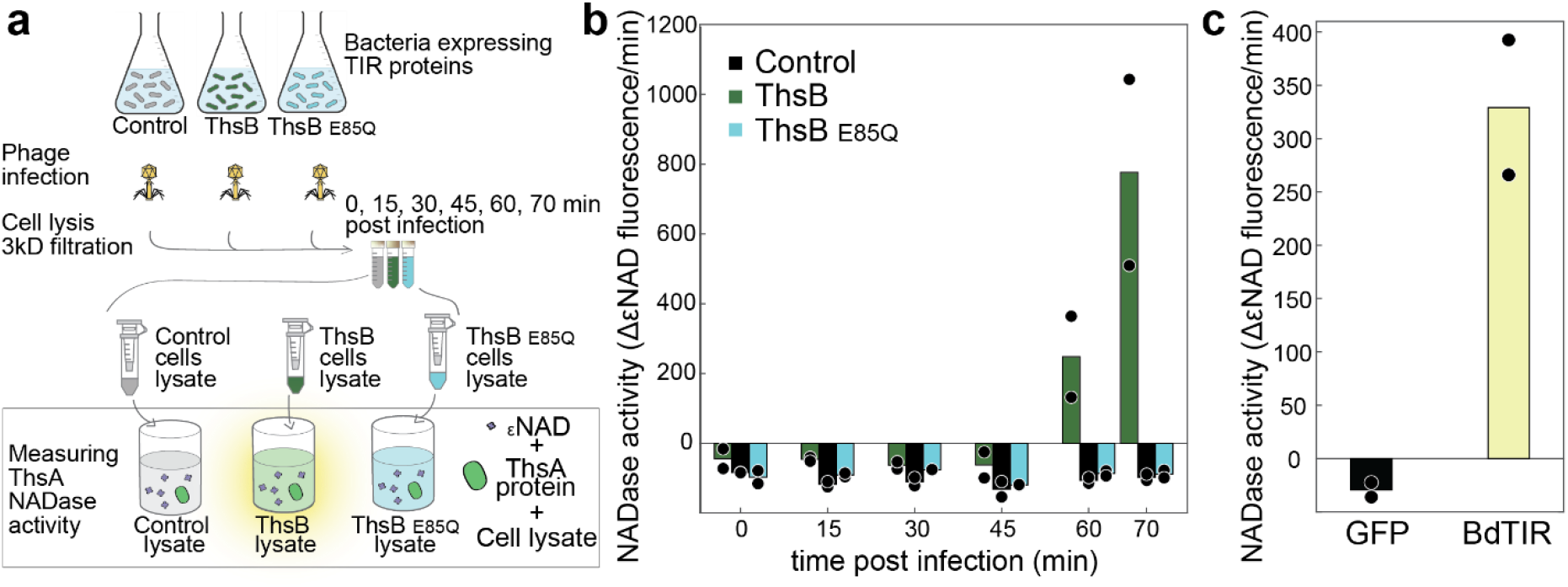
The NADase activity of ThsA is activated by small molecules generated in TIR-expressing infected cells. **(a)** Schematic representation of the experiment. Cells expressing either ThsB protein, ThsB^E85Q^ mutant or GFP (control) were sampled before infection (time 0) and at 15, 30, 45, 60 and 70 minutes post infection with SPO1 at an MOI of 5. Sampled cells were then pelleted, lysed and filtered through a 3 kDa filter to retain only the small molecule fraction. Purified ThsA protein was then incubated with the lysates and its NADase activity was measured using a nicotinamide 1,N6-ethenoadenine dinucleotide (εNAD) cleavage fluorescence assay. **(b)** NADase activity of purified ThsA protein incubated with lysates derived from control cells (black), ThsB-expressing cells (green) or cells expressing the mutant ThsB_E85Q_ (cyan), during infection. NADase activity was calculated as the rate of change in εNAD fluorescence during the linear phase of the reaction. Negative values represent bleaching of background fluorescence. Bars represent the mean of two experiments, with individual data points overlaid. Raw fluorescence measurements are presented in Figure S2. **(c)** NADase activity of ThsA when incubated with lysates derived from *E. coli* cells overexpressing GFP control (black) or BdTIR (yellow).

To inquire if the bacterial TIR-generated molecular agent that activates ThsA is similar to the molecules generated by plant TIRs, we used lysates from *Escherichia coli* cells overexpressing a TIR-domain protein from the plant *Brachypodium distachyon* (BdTIR), which was shown to constitutively produce v-cADPR when expressed in *E. coli*^3^. Lysates from BdTIR-expressing *E. coli* were able to trigger the NADase activity of ThsA (Figure 2C). In contrast, ThsA was not active in the presence of the canonical cADPR molecule, even when cADPR was supplied at millimolar concentration (Figure S2B). These results suggest that the bacterial Thoeris TIR and the plant BdTIR are functionally similar, and that secondary messenger molecules produced by both can activate the Thoeris effector gene.

### The TIR-produced signaling molecule is an isomer of cADPR

To further study the signaling molecule produced by the Thoeris TIR protein, we used liquid chromatography followed by untargeted mass spectrometry (LC-MS) to analyze the metabolite content in *B. subtilis* cells expressing Thoeris during infection by phage SPO1. These experiments were performed using a Thoeris system in which ThsA was mutated in its NADase active site, to prevent premature cell death and allow the accumulation of the signaling molecule to detectable levels prior to cell death. We observed a unique molecule that became detectable in the TIR-expressing infected cells 60 minutes after infection (Figure 3A). The molecule was not detected in uninfected cells or at earlier time points post infection, and was also absent from infected cells that expressed the mutated TIR protein ThsBE85Q (Figure 3A, S3A). These results suggest that this unique molecule is generated by the TIR protein in response to phage infection. The m/z value of the identified molecule was 542.0683 (positive ionization mode), which is within the expected measurement error of the predicted protonated mass of cyclic ADP-ribose (cADPR, m/z = 542.0684). However, the unique molecule eluted from the LC column at a different retention time than the canonical cADPR standard, indicating that it is structurally distinct from the canonical cADPR (Figure 3B). Tandem MS fragmentation analysis (MS/MS) demonstrated that the Thoeris-produced molecule generates molecular fragments similar to those generated by the canonical cADPR (Figure 3C), further supporting the hypothesis that the Thoeris TIR-produced molecule is an isomer of cADPR.

**Figure 3.**
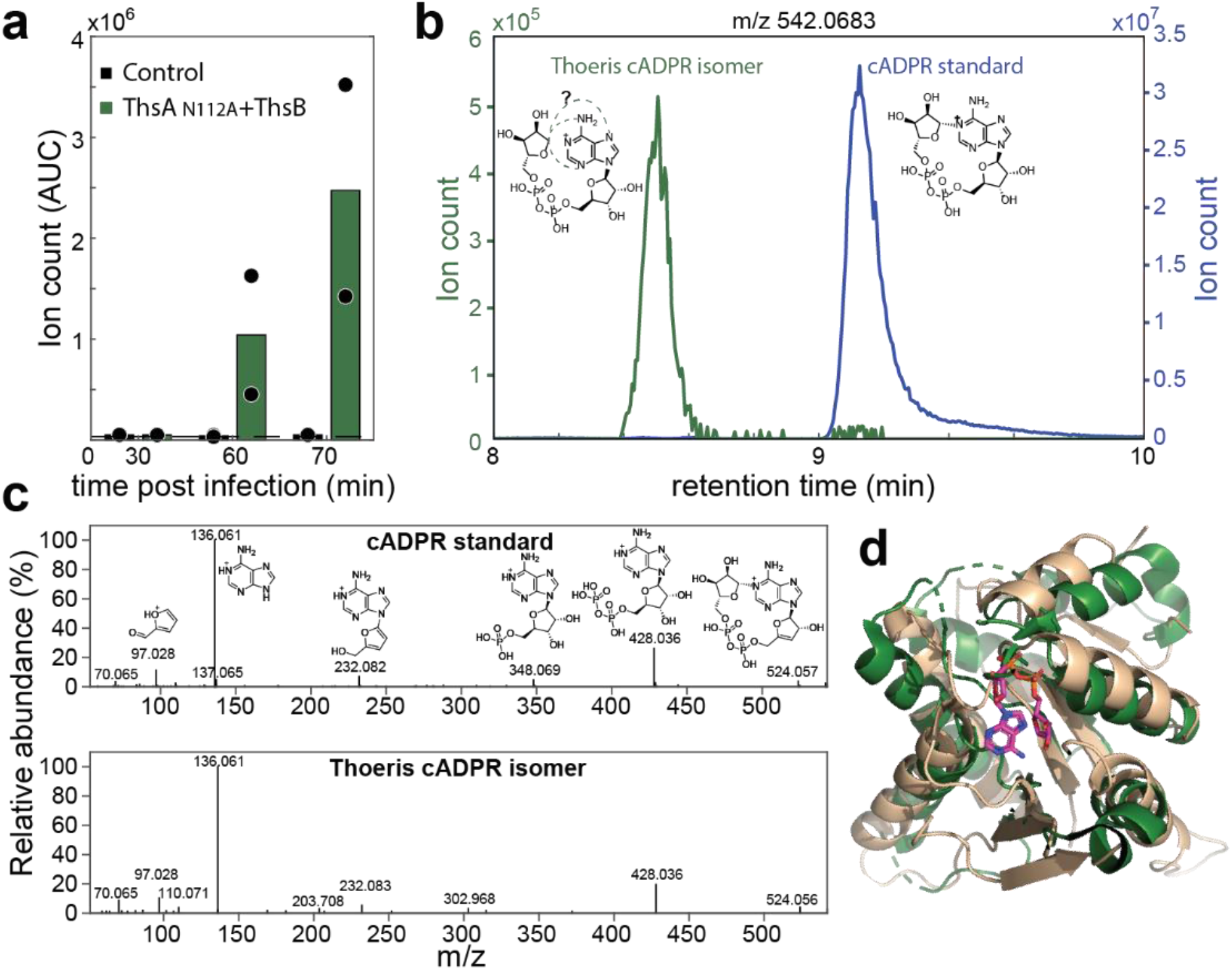
ThsB produces an isomer of cADPR upon phage infection. **(a)** A unique molecule with an m/z value of 542.0683 and retention time of 8.5 minutes appears in infected cells that express wild type ThsB and mutated ThsA_N112A_. Cells were infected with SPO1 at MOI of 5, lysates were taken from several time points post infection, filtered through a 3 kDa filter, and analyzed by LC-MS in positive ionization mode. Bars represent the mean of two experiments, with individual data points overlaid. Dashed line represents limit of detection. **(b)** Chromatograms of the peak measured in panel **a**, detected in lysates of Thoeris-expressing cells 70 minutes post infection (green curve, green Y axis) and of standard cADPR (blue curve, blue Y axis), demonstrating retention time difference. The molecular structure of standard cADPR and possible cyclization bonds in the Thoeris cADPR isomer are presented. **(c)** MS/MS fragmentation spectra of standard cADPR (top) and the Thoeris-derived cADPR isomer (bottom). Hypothesized structures of MS/MS fragments of cADPR are presented. **(d)** Structural superimposition of the SLOG domain of ThsA (PDB ID 6LHX, green)^11^ and the SLOG domain of human TRPM2 bound to ADPR (PDB ID 6PUS, wheat)^14^.

In addition to the SIR2 NADase domain at its N-terminus, ThsA contains another domain, at its C-terminus, called SLOG^11,15,16^. SLOG domains were hypothesized to bind nucleotide-derived signaling molecules, and specifically NAD derivatives such as ADPR molecules^15,16^. Domains homologous to SLOG are found in human cation channels of the TRPM family^15^ which are triggered by ADPR and its derivatives^17,18^, and it was shown that the ADPR molecule directly binds to the SLOG domain of human TRPM2 to activate it^14^. Structural superimposition of the SLOG domain of ThsA and the ADPR-bound SLOG domain of TRPM2 demonstrate their structural homology in the ADPR-binding pocket region (Figure 3D). We therefore hypothesize that ThsA is activated when the TIR-produced cADPR isomer binds to its C-terminal SLOG domain.

### TIR-domain proteins determine the specificity of Thoeris against phages

In natural Thoeris systems, the *thsA* SIR2-domain gene is often accompanied by multiple *thsB* TIR-domain genes^8^ (Figure 4A). Different *thsB* genes at the same locus are usually divergent from one another, often sharing very little sequence identity beyond the general structure of the TIR domain and its conserved active site^8,16^. We hypothesized that diverse ThsB proteins within the same locus could be responsible for recognition of different phage-associated molecular patterns, akin to the roles of TIR-domain proteins in eukaryotic immune systems. Under this hypothesis, each TIR-domain protein would serve to recognize a different phage infection marker. Once this marker had been identified, the TIR domain would produce the cADPR isomer molecule to activate the effector ThsA NADase.

**Figure 4.**
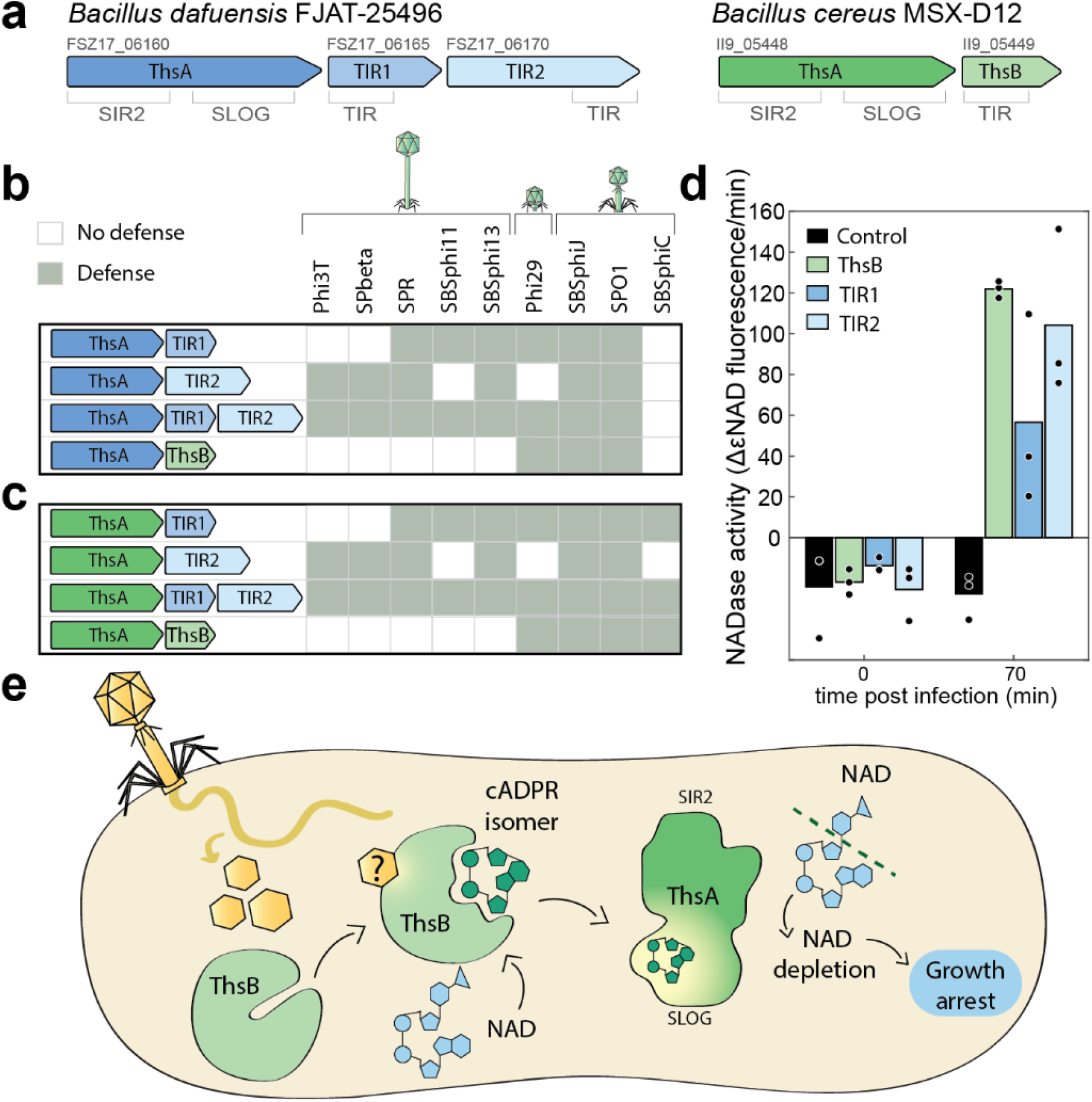
Thoeris TIR proteins determine defense specificity. **(a)** Schematic representation of the Thoeris systems from *Bacillus dafuensis* FJAT-25496 (blue) and *B. cereus* MSX-D12 (green). Gene locus tags and protein domains are indicated. **(b)** Defense patterns of Thoeris systems containing combinations of ThsA from *B. dafuensis* and ThsB proteins from *B. dafuensis* or *B. cereus*. In each combination, ThsA was induced with 0.2% xylose and ThsB with 100μM IPTG. Reduction of more than 10-fold in the efficiency of plating (EOP) on system-containing cells as compared to control cells is marked by a grey rectangle. EOP data used to generate this figure were collected in three independent replicates, and are presented in Figure S4A. **(c)** Same as panel b, for combinations of ThsA from *B. cereus* and ThsB proteins from *B. dafuensis* or *B. cereus*. **(d)** NADase activity of purified ThsA protein from *B. cereus*, incubated with lysates derived from control cells (black), or cells expressing TIR-domain proteins of *B. cereus* (green) and *B. dafuensis* (blue shades). Lysates were collected from cells infected by SPO1, at an MOI of 5, prior to infection (time 0) or 70 minutes post infection. NADase activity was calculated as the rate of change in εNAD fluorescence during the linear phase of the reaction. Bars represent the mean of three experiments, with individual data points overlaid. **(e)** A model for the mechanism of action of Thoeris. Phage infection is sensed by the ThsB TIR-domain protein, which becomes enzymatically active to produce the cADPR isomer signaling molecule. The molecule is sensed by the ThsA SLOG domain, activating the enzymatic activity of the SIR2 domain to deplete the cell of NAD and lead to abortive infection.

To test this hypothesis, we examined the Thoeris locus found in *Bacillus dafuensis* FJAT-25496, which includes two TIR-domain *thsB* genes present consecutively on the same operon with a single upstream *thsA* gene (Figure 4A). The two TIR proteins of this system have no identified homology when aligned using BLASTP^19^, but both contain a TIR domain identifiable by the HHpred tool^20^. We constructed *B. subtilis* strains expressing the *B. dafuensis* ThsA together with either of the two ThsB proteins (called here TIR1 and TIR2) or with both proteins together (Figure 4A-B). *B. subtilis* cells expressing ThsA+TIR1 were protected from infection by a set of phages including phi29 and SBSphi11, but were sensitive to phages phi3T and SPBeta (Figure 4B, Figure S4A). In contrast, cells expressing ThsA+TIR2 were protected from phages phi3T and SPBeta, but sensitive to phages phi29 and SBSphi11 (Figure 4B, Figure S4A). Cells expressing ThsA+TIR1+TIR2 were protected from all four phages, showing that the defense range conferred by the individual TIR proteins is additive (Figure 4B, Figure S4A). These results are in line with the hypothesis that each TIR-containing ThsB protein recognizes a different phage infection marker to initiate Thoeris defense. Notably, both ThsA+TIR1 and ThsA+TIR2 constructs provided defense against phages SPO1, SBSphiJ, SPR and SBSphi13, implying that these phages present infection markers that activate both TIRs.

Interestingly, a hybrid system in which we co-expressed the ThsA of *B. dafuensis* together with the ThsB TIR-containing protein from *B*. cereus was also functional in defense against phages, indicating that the *B. dafuensis* ThsA responds to the same molecular signal (cADPR isomer) produced by the *B. cereus* TIR protein (Figures 4B, S4A). In further support of this, TIR proteins from *B. dafuensis* were functional in defense when co-expressed with *B. cereus* ThsA (Figures 4C, S4A). Moreover, filtered lysates from SPO1-infected cells expressing each of the two TIR proteins from *B. dafuensis* were able to trigger the NADase activity of ThsA from *B. cereus in vitro*, indicating that both these TIR-domain proteins produce the ThsA-activating cADPR isomer when triggered by phage infection (Figure 4D).

Cloning of the various TIR proteins with a ThsA_N112A_ mutant did not provide defense against any of the phages, demonstrating that an intact ThsA is essential for defense regardless of the associated TIR-domain protein (Figure S4B). With the exception of SBSphiC, the defense specificity against all phages seemed to be determined by the identity of the TIR-domain protein, rather than the ThsA protein, further supporting the role of the TIR-domain proteins as the specificity determinants of Thoeris (Figure 4B-C). Defense against SBSphiC was only seen when the ThsA from *B. cereus* was included (Figure 4B-C); this could be explained by a specific ability of SBSphiC to counteract the activity of *B. dafuensis* ThsA via a mechanism that does not prevent the activation of *B. cereus* ThsA.

## Discussion

Combined together, our results point to a model for the mechanism of the Thoeris anti-phage defense system (Figure 4E). The TIR-domain protein ThsB is responsible for recognizing phage infection. Once infection has been detected, the TIR domain becomes enzymatically active and catalyzes the production of the cADPR isomer molecule. This molecule, in turn, serves as a signaling agent that binds the ThsA effector, most probably via its C-terminal SLOG domain, and activates its NADase activity. The NADase effector then depletes NAD from the cell, leading to cellular conditions that cannot support phage replication and presumably lead to cell death (Figure 4E).

Our data show that TIR domains in the Thoeris anti phage defense systems produce a signaling molecule that activates the associated effector to abort phage infection. Similar molecules produced by plant TIR domains were recently hypothesized to serve as secondary messengers for plant immunity, but their exact roles in plants and their putative receptors are currently unknown^6,7^. Our results verify that TIR-derived cADPR isomers can serve as immune secondary messengers, and can inform future studies in plant model systems to reveal the role of such TIR-produced molecules in the plant immune response to pathogens.

The Thoeris system is functionally analogous to a recently discovered family of anti-phage defense systems called CBASS (cyclic oligonucleotide-based antiphage signaling system), which are also abortive infection systems that rely on secondary messenger intracellular signaling^12,21–25^. Structural and functional data show that the CBASS system is the evolutionary ancestor of the animal cGAS-STING antiviral pathway, which senses viral infection and relies on cyclic GMP-AMP signaling to activate immunity^12^. We find it remarkable that two prokaryotic defense systems that are based on small molecule signaling, CBASS and Thoeris, have evolutionarily contributed core functions to the animal and plant immune systems, respectively.

The exact molecular structure of the signaling cADPR isomer remains to be determined. Nucleotide-based immune signaling molecules in bacteria were shown to function in nanomolar or low micromolar concentrations^12,23,25^, limiting analytical studies. Indeed, while the v-cADPR molecule was observed in multiple studies of plant TIR domains^3,4,7^, its molecular structure has not yet been deciphered. It was hypothesized that this molecule may be a cyclization isomer of cADPR, with the cyclic bond between the ribose and the adenine formed via a different position on the adenine base^26^. Alternatively, the signaling molecule could be a stereoisomer of cADPR, or a non-cyclical ADP-ribose variant that shares the same mass with cADPR. Our data show that a molecule produced when the plant BdTIR is expressed in *E. coli* functionally activates Thoeris ThsA. However, determining whether the bacterial and plant signaling molecules are identical awaits deciphering of the exact structures of these molecules.

Our data show that ThsA activity depletes NAD from infected cells leading to abortive infection. NAD depletion was previously shown to be toxic to bacterial cells^12,27–29^, and was recently demonstrated in other abortive infection systems^12^. However, we cannot rule out the possibility that NAD depletion alone is not the direct cause for the observed cell death. For example, the SIR2 domain of ThsA could be using large amounts of NAD as a cofactor for another enzymatic activity that causes the observed abortive infection.

TIR domains have been canonically described as protein-protein interaction platforms that mediate immune signaling of Toll-like receptors in animals^1^. Observations from recent years, including those described in the current study, show that TIRs can also function as enzymes. TIR domains were shown to serve as NAD depletion enzymes as in the case of the human SARM1 protein^30^, TIR-containing effector proteins in pathogenic bacteria^26,31^, and TIR-STING proteins in bacterial CBASS systems^12^. Moreover, in plants^3,4^, and as shown here in bacteria, TIR domains can utilize NAD for the production of signaling molecules, and we now demonstrate that these signals activate downstream components of the Thoeris system to trigger cell death. Our data imply that TIR-based antiviral immune signaling in bacteria may have been the ancestral form of plant TIR-containing innate immunity mechanisms.

## Acknowledgements

We thank the Sorek laboratory members, Maya Voichek, Asaf Levy, Daniel Dar, Virginijus Šikšnys, and Mindaugas Zaremba for comments on earlier versions of this manuscript. We thank Yinon M. Bar-On for useful discussion during the project, Carmel Avraham and Taya Fedorenko for their assistance with the experiments, and Aude Bernheim for her continuous advice and support throughout this project. R.S. was supported, in part, by the European Research Council (grant ERC-CoG 681203), the Ernest and Bonnie Beutler Research Program of Excellence in Genomic Medicine, the Minerva Foundation with funding from the Federal German Ministry for Education and Research, the Knell Family Center for Microbiology, and the Yotam project and the Weizmann Institute Sustainability And Energy Research (SAERI) initiative. G.O. was supported by the SAERI doctoral fellowship. A.M. was supported by a fellowship from the Ariane de Rothschild Women Doctoral Program and, in part, by the Israeli Council for Higher Education via the Weizmann Data Science Research Center, and by a research grant from Madame Olga Klein-Astrachan.

## Materials and methods

### Plasmid construction for *Bacillus* shuttle vectors

Accession numbers of the genes used in this study are listed in Table 1. Gene ORFs were synthesized by Genscript Corp. *Bacillus cereus* MSX-D12 *thsA* gene included 6 additional bases upstream to the ORF annotated in NCBI (GenBank accession: EJR09240.1) to include an upstream ATG. The *thsA* gene of *B. dafuensis* FJAT-25496 included 27 additional bases upstream to the annotated ORF to include a possible alternative upstream ATG. Synthesized DNA was used as a template for PCR for subsequent cloning into shuttle vectors for *Bacillus* transformation. All PCR reactions were performed using KAPA HiFi HotStart ReadyMix (Roche cat # KK2601). Plasmid maps are attached as Supplementary Files 1-9. All primers used are listed in Table 2.

**Table 1.** Genes cloned in this study

| Genome | Gene | GenBank accession | Refseq accession |
| --- | --- | --- | --- |
| *Bacillus cereus* MSX-D12 | *thsA* | EJR09240.1 | WP_002078322.1 |
| *Bacillus cereus* MSX-D12 | *thsB* | EJR09241.1 | WP_001129548.1 |
| *Bacillus dafuensis* FJAT-25496 | *thsA* | QED46885.1 | WP_057775117.1 |
| *Bacillus dafuensis* FJAT-25496 | *TIR1* | QED46886.1 | WP_057775115.1 |
| *Bacillus dafuensis* FJAT-25496 | *TIR2* | QED46887.1 | WP_057775113.1 |

**Table 2.**
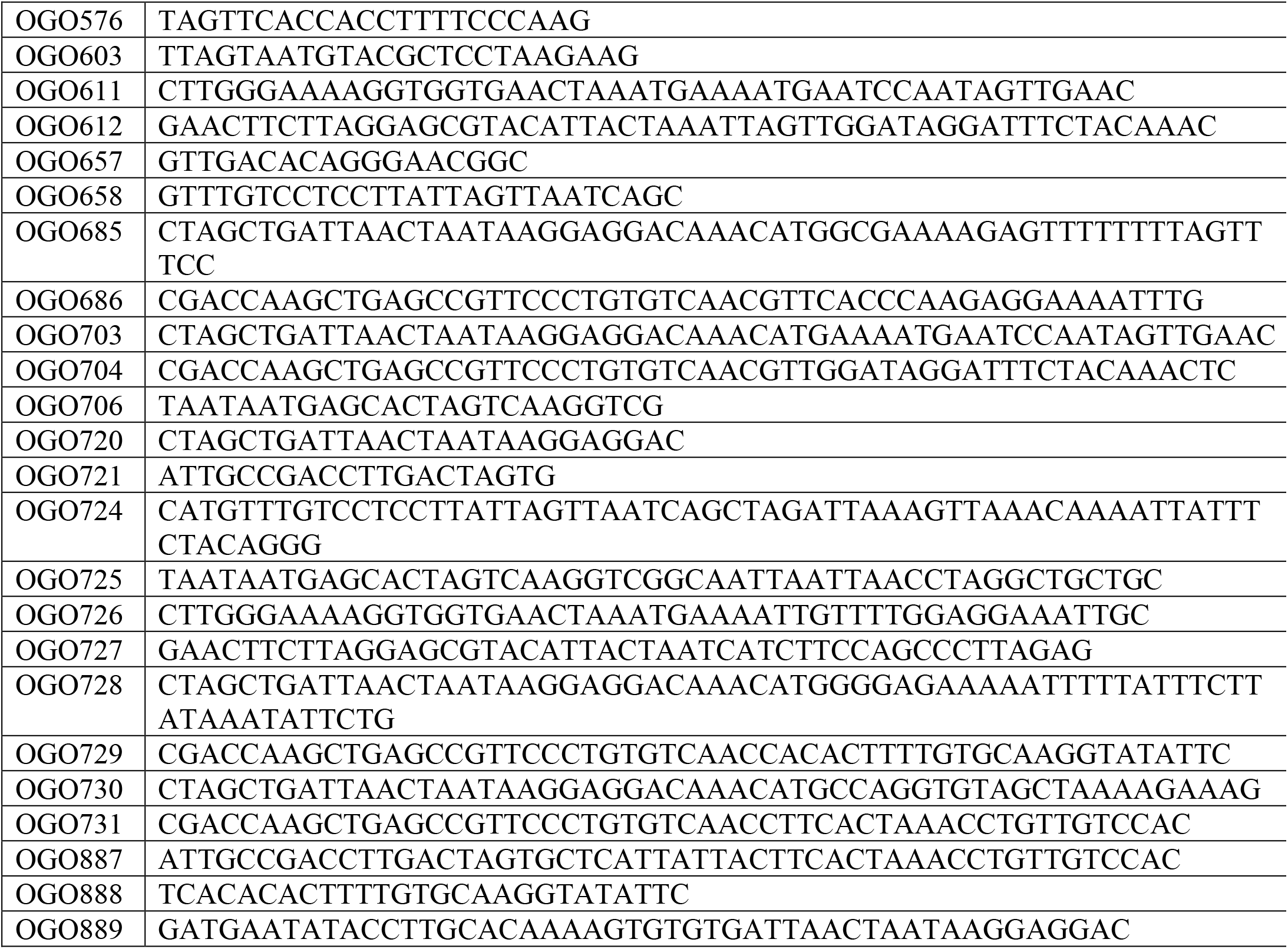
Primers used in this study

The *thsA* gene of *Bacillus cereus* MSX-D12 was amplified using primers OGO611+OGO612. *Bacillus dafuensis* FJAT-25496 *thsA* gene was amplified using primers OGO726+OGO727. The *thsA* genes were cloned into a shuttle vector under the control of Pxyl promoter (Supplementary Files 1-2). The shuttle vector contained chloramphenicol resistance and a *thrC* integration cassette for genomic integration and propagation in *Bacillus subtilis*. The vector backbone was amplified using primers OGO576+OGO603.

The *thsB* genes were cloned into a second shuttle vector under the control of a Phspank promoter, and fused to a C-terminal TwinStrep tag (Supplementary Files 3-5). The shuttle vector contained spectinomycin resistance and an *amyE* integration cassette for genomic integration and propagation in *Bacillus subtilis*. For the control strain, superfolder GFP gene (sfGFP) was cloned instead of the *thsB* genes, and the *thrC* locus was uninterrupted (Supplementary File 6). *Bacillus cereus* MSX-D12 *thsB* was amplified using primers OGO685+OGO686. *Bacillus dafuensis* FJAT-25496 *TIR1* was amplified using primers OGO728+OGO729. *Bacillus dafuensis* FJAT-25496 *TIR2* was amplified using primers OGO730+OGO731. The vector backbone was amplified using primers OGO657+OGO658.

The double TIR construct of *Bacillus dafuensis* FJAT-25496 (TIR1+TIR2) was constructed by PCR amplification of the TIR2 gene and its ribosome binding site from its shuttle vector and cloning it downstream to the TIR1 gene on the *amyE* shuttle vector, to create a bicistronic transcript expressed from the Phspank promoter (Supplementary File 7). The TIR1 vector backbone was amplified using primers OGO706+OGO888, and the TIR2 insert was amplified using primers OGO887+OGO889.

Cloning was performed using NEBuilder HiFi DNA Assembly kit (NEB, cat # E2621). Both shuttle vectors were propagated in *E. coli* DH5a using a p15a origin of replication with 100 μg/ml ampicillin selection. Plasmids were miniprepped from *E. coli* DH5a prior to transformation into *B. subtilis* BEST7003^32^.

Mutagenesis of *Bacillus cereus* MSX-D12 ThsB E85Q (GAA>CAA) was carried out by Genscript Corp. Mutagenesis of *Bacillus cereus* MSX-D12 ThsA N112A was previously described^8^.

### Bacillus transformation

Transformation to *B. subtilis* BEST7003 was performed using MC medium as previously described^33^. MC medium was composed of 80 mM K2HPO4, 30 mM KH_2_PO_4_, 2% glucose, 30 mM trisodium citrate, 22 μg/ml ferric ammonium citrate, 0.1% casein hydrolysate (CAA), 0.2% potassium glutamate. From an overnight starter of bacteria, 10 μl were diluted in 1 ml of MC medium supplemented with 10 μl 1M MgSO_4_. After 3 hours of incubation (37°C, 200 rpm), 300 μl of the culture was transferred to a new 15 ml tube and ~200 ng of plasmid DNA was added. The tube was incubated for another 3 hours (37°C, 200 rpm), and the entire reaction was plated on Lysogeny Broth (LB) agar plates supplemented with 5 μg/ml chloramphenicol or 100 μg/ml spectinomycin and incubated overnight at 30°C.

### Phage cultivation and plaque assays

Phages were previously received from the Bacillus Genetic Stock Center (BGSC). BGSC IDs for the phages used are 1P4 for SPO1, 1L1 for phi3T, 1L5 for SPβ, 1L56 for SPR, 1P7 for SPP1. Phage phi29 was received from DSMZ (DSM 5546). Phages SBSphiC and SBSphiJ were previously isolated by us^8^ and are available from BGSC (BGSCID 1P46 and 1P47 respectively). Phages SBSphi11 and SBSphi13 were isolated by us during this study from soil samples as previously described^8^.

Phages were propagated by infection of exponentially growing *B. subtilis* BEST7003 culture in magnesium-manganese broth (MMB) (LB + 0.1 mM MnCl_2_ + 5 mM MgCl_2_) at MOI of 1:100-1:1000, incubated at 37°C with shaking (200 rpm) until clearing of the culture, the lysate was centrifuged and the supernatant was filter sterilized through 0.45 μm filter.

Phage titer was determined using the small drop plaque assay method^34^. 300 μl of overnight culture of bacteria were mixed with 25 ml MMB+0.5% agar, poured in a 10 cm square plate and incubated for 1 hour at room temperature. For cells that contained inducible constructs, the inducers were added to the agar at a concentration of 0.2% xylose and 5 μM IPTG for experiments presented in Figure S1, or 0.2% xylose and 100 μM IPTG for experiments presented in Figure 4 and Figure S4. The phage stock was tenfold serially diluted and 10 μl drops of diluted phage lysate were placed on the solidified agar. After the drops have dried up, the plates were inverted and incubated at room temperature overnight, and plaques were counted to determine phage titer. Phage efficiency of plating (EOP), representing the magnitude of defense, was determined as the ratio between the number of plaques on the control strain expressing GFP and the number of plaques on the relevant Thoeris strain. When individual plaques could not be counted due to small size, a faint lysis zone across the drop area was considered to be 100 plaques.

### Infection dynamics in liquid culture

Non-induced overnight cultures of *B. subtilis* cells were diluted 1:100 into fresh MMB medium supplemented with 0.2% xylose and 5 μM IPTG. Cultures were incubated at 37°C with shaking (200 rpm). When cells reached OD_600_ of 0.3, 180 μl were dispensed into each microwell in a 96-well plate and infected with 20 μl of SPO1 phage lysate to the desired MOI. Plates were immediately transferred to a plate reader (Tecan Infinite 200) set to 25°C with 4.5 mm orbital shaking and OD_600_ monitored every 10 minutes.

### NAD levels measurements *in vivo*

Overnight cultures were diluted 1:50 in 100ml MMB supplemented with 0.2% xylose and 5 μM IPTG and grown at 37°C, 200 rpm shaking, until OD_600_ of 0.3. A sample of 15 ml for uninfected culture (time 0) was then removed, and SPO1 phage stock was added to the culture to reach MOI of 5. Flasks were incubated at 25°C, 200 rpm shaking, for the duration of the experiment. 15 ml samples were then collected at times 15, 30, 45, 60, 70 min post infection. Immediately upon sample removal the sample tube was placed in ice and centrifuged at 4°C for 8 minutes to pellet the cells. The supernatant was discarded and the tube was frozen at - 80°C. To extract the metabolites, 600 μl of 100mM phosphate buffer supplemented with 4 mg/ml lysozyme (Sigma cat # L6876) was added to each pellet. Tubes were then incubated for 5 minutes at 37°C, and returned to ice. The thawed sample was transferred to a FastPrep Lysing Matrix B in a 2 ml tube (MP Biomedicals cat # 116911100) and lysed using FastPrep bead beater for 40 seconds at 6 m/s. Tubes were then centrifuged at 4°C for 15 min at 15,000 *g*. Supernatant was transferred to Amicon Ultra-0.5 Centrifugal Filter Unit 3 KDa (Merck Millipore cat # UFC500396) and centrifuged for 45 min at 4°C at 12,000 *g*. Filtrates were taken for LC-MS analysis.

Profiling of polar metabolites was done as previously described^35^ with minor modifications as described below. Briefly, the analysis was performed using an Acquity I class UPLC System combined with a mass spectrometer (Thermo Exactive Plus Orbitrap) which was operated in a positive ionization mode using a mass range 300-700 m/z. The LC separation was done using the SeQuant Zic-pHilic (150 mm × 2.1 mm) with the SeQuant guard column (20 mm × 2.1 mm) (Merck). The Mobile phase B was acetonitrile and Mobile phase A was 20 mM ammonium carbonate plus 0.1% ammonia hydroxide in water. The flow rate was kept at 200 μl/min and the gradient was as follows: 75% B (0-2 minutes), decrease to 25% B (2-14 minutes), 25% B (14-18 minutes), increase back to 75% B (18-19 minutes), 75% B (19-23 minutes). NAD and ADPR peaks were identified in the data using a synthetic standard run (Sigma-Aldrich N8285 and A0752, respectively), and area under the peak was quantified using MZmine 2.53^36,37^ with an accepted deviation of 5ppm.

### Cloning of BdTIR

The BdTIR gene from *Brachypodium distachyon* Bd21 (GenBank accession KQK15683.1; Refseq accession XP_003560074.3) was codon-optimized for *E. coli* expression using the Genscript codon optimization tool, and synthesized with an N-terminal TwinStrep tag as described in Wan et al^3^. The gene was synthesized and cloned by Genscript Corp. into a pET30a vector between the NdeI and XhoI sites of the plasmid to fuse a C-terminal 6XHis tag from the plasmid backbone (Supplementary File 8). Plasmids were then electroporated into *E. coli* BL21(DE3) cells, and grown in LB medium supplemented with 50 μg/ml kanamycin.

### ThsA protein purification

Plasmid pAB151^38^ was used to generate a protein expression vector. The plasmid backbone was amplified using primers OGO724+OGO725, and assembled with the sfGFP-TwinStrep segments from the *amyE* shuttle vector amplified using primers OGO720+OGO721. The *Bacillus cereus* MSX-D12 *thsA* gene was amplified from the *thrC* shuttle vector using primers OGO703+OGO704 and assembled into the *amyE* shuttle vector backbone amplified using primers OGO657+OGO658 to be fused with the C-terminal TwinStrep tag. Then, the tagged gene was amplified using primers OGO720+OGO721 and cloned into pAB151 backbone (Supplementary File 9). Plasmids were constructed using NEBbuilder HiFi DNA Assembly kit, propagated in *E. coli* DH5a cells, miniprepped, and then electroporated into *E. coli* BL21(DE3) cells. Cells were grown in LB supplemented with 30 μg/ml chloramphenicol.

For protein expression and purification, an overnight culture of *E. coli* BL21(DE3) was diluted 1:100 into fresh LB, grown to OD_600_ of 0.6, and induced using 100 μM IPTG. The culture was then grown at 20°C overnight. Cells were pelleted by centrifugation in 50ml culture aliquots, and frozen at −80°C. To purify the protein, 1 ml of Strep wash buffer (IBA cat # 2-1003-100) supplemented with 4 mg/ml lysozyme was added to each pellet and incubated for 10 minutes at 37°C with shaking until thawed and resuspended. Tubes were then transferred to ice, and the resuspended cells transferred to a FastPrep Lysing Matrix B in 2 ml tube (MP Biomedicals cat # 116911100). Samples were lysed using FastPrep bead beater for 40 seconds at 6 m/s. Tubes were centrifuged for 15 minutes at 15,000 *g*. Per each pellet, 30 μl of MagStrep “Type 3” XT beads (IBA cat # 2-4090-002) were washed twice in 300 μl wash buffer, and the lysed cell supernatant was mixed with the beads and incubated for 30-60 minutes, rotating in 4°C. The beads were then pelleted on a magnet, washed twice with wash buffer, and purified protein was eluted from the beads in 10 μl of BXT elution buffer (IBA cat # 2-1042-025). Purified ThsA protein was freshly prepared for each experiment.

### Cell lysate preparation

Overnight cultures of the ThsB-expressing, ThsA_N112A_+ThsB-expressing, and GFP-expressing (control) strains were diluted 1:100 in 250 ml MMB supplemented with 0.2% xylose and 100 μM IPTG, and incubated at 37°C with shaking (200 rpm) until reaching OD_600_ of 0.3. SPO1 phage stock was added to the culture to reach an MOI of 5. Flasks were incubated at 25°C with shaking (200 rpm) for the duration of the experiment. 50 ml samples were collected at various time points post infection. Immediately upon sample removal the sample tube was placed in ice, and then centrifuged at 4°C for 8 minutes to pellet the cells. The supernatant was discarded and the tube was frozen at −80°C. To extract the metabolites, 600 μl of 100mM phosphate buffer at pH 8, supplemented with 4 mg/ml lysozyme, was added to each pellet. The tubes were then incubated for 5 minutes at 37°C, and returned to ice. The thawed sample was transferred to a FastPrep Lysing Matrix B 2 ml tube (MP Biomedicals cat # 116911100) and lysed using a FastPrep bead beater for 40 seconds at 6 m/s, two cycles. Tubes were then centrifuged at 4°C for 15 minutes at 15,000 *g*. Supernatant was transferred to Amicon Ultra-0.5 Centrifugal Filter Unit 3 kDa (Merck Millipore cat # UFC500396) and centrifuged for 45 minutes at 4°C at 12,000 *g*. Filtrate was taken and used for LC-MS analysis or enzymatic activity assays. For the experiment presented in Figure 3A, 2 pellets from two 50 ml culture samples were united prior to bead beating to increase metabolite concentrations in the sample used for LC-MS detection of Thoeris cADPR isomer.

For BdTIR lysate preparation, an overnight culture of *E. coli* BL21(DE3) cells carrying the BdTIR plasmid was diluted 1:100 in LB, incubated at 37°C with shaking (200 rpm) for 3 hours, induced with 100 μM IPTG, and incubated for 3 additional hours. The culture was then collected in 50 ml tubes, centrifuged for 8 minutes at 4°C to pellet the cells, the supernatant was discarded and pellets were frozen at −80°C. Cell lysate preparation was performed as described above. For the control lysates, the same process was done with cells expressing GFP from the pAB151 vector backbone.

### ThsA NADase activity assay

Purified ThsA protein was freshly prepared and protein concentration was quantified using Qubit Protein Assay Kit (ThermoFisher cat # Q33212). Protein was diluted in wash buffer to a concentration of 147 ng/μl. In each microwell of black 96-well half area plates (Corning cat # 3694), 43 μl of cell lysate or 100 mM phosphate buffer was placed, and the plate was kept on ice. 2 μl of the diluted protein was then added to each plate for a final concentration of 100 nM protein in the final 50 μl reaction. 5 μl of 5 mM Nicotinamide 1,N6-ethenoadenine dinucleotide (εNAD, Sigma cat # N2630) solution were added to each well immediately prior to the beginning of measurements and mixed by pipetting to reach a concentration of 500 μM in the 50 μl final volume reaction. Plates were incubated inside a Tecan Infinite 200 plate reader at 25°C, and measurements were taken every 1 minute at 320 nm excitation wavelength and 400 nm emission wavelength.

### LC-MS identification of Thoeris cADPR isomer

Cell lysates were prepared as described above and sent for analysis at MS-Omics Ltd. Sample analysis was carried out by MS-Omics as follows. Samples were diluted 1:1 in 10% ultra-pure water and 90% acetonitrile containing 10 mM ammonium acetate at pH 9, then filtered through a Costar^®^ Spin-X ^®^ centrifuge tube filter 0.22 μm nylon membrane. The analysis was carried out using a Vanquish™ Horizon UHPLC System coupled to Q Exactive™ HF Hybrid Quadrupole-Orbitrap™ Mass Spectrometer (Thermo Fisher Scientific, US). The UHPLC was performed using an Infinity Lab PoroShell 120 HILIC-Z PEEK lined column with the dimension of 2.1 x 150mm and particle size of 2.7μm (Agilent Technologies). The composition of Mobile phase A was 10 mM ammonium acetate at pH 9 in 90% Acetonitrile LC-MS grade (VWR Chemicals, Leuven) and 10 % Ultra-pure water from Direct-Q^®^ 3 UV Water Purification System with LC-Pak^®^ Polisher (Merck KGaA, Darmstadt) and mobile phase B was 10 mM ammonium acetate at pH 9 in ultra-pure water with 15 μM medronic acid (InfinityLab Deactivator additive, Agilent Technologies). The flow rate kept at 250 μl/ml consisting of a 2 minutes hold at 10% B, increased to 40% B at 14 minutes, held till 15 minutes, decreased to 10% B at 16 minutes and held for 8 minutes. The column temperature was set at 30 °C and an injection volume of 5 μl. An electrospray ionization interface was used as ionization source. Analysis was performed in positive ionization mode from m/z 300 to 900 at a mass resolution of 120000 (at m/z 200). Peak areas were extracted using Compound Discoverer 3.1 (Thermo Fisher Scientific, US) with an accepted deviation of 3 ppm. The standard cADPR peak was identified using a synthetic standard run (Sigma-Aldrich, C7344). MS/MS spectra collection was performed using the same instrument at a resolution of 30,000. Fragmentation was done through a higher-energy collisional dissociation cell using a normalized collision energy of 20, 40 and 60 eV where the spectrum is the sum of each collision energy.

### Structural superimposition of ThsA and TRPM2

The structures of *B. cereus* MSX-D12 ThsA (PDB ID 6LHX) and human TRPM2 bound to Ca^+2^ and ADPR (PDB ID 6PUS) were downloaded from PDB^39^. The SLOG domain of ThsA was defined as residues 281-471 of chain A of 6LHX. The SLOG domain of TRPM2 was defines as residues 167-437 of chain C of 6PUS. The domains were aligned using the “super” structural imposition function of PyMOL (Schrödinger LLC) version 2.4.0 with 25 cycles limit. Residues 247-277 and 370-412 of 6PUS did not align with ThsA and were not represented in the visualization shown in Figure 3D for clarity.

## Supplementary Materials

### Supplementary files

**Supplementary File 1**. Plasmid map of pGO1_thrC_Pxyl_cereus_ThsA, shuttle vector for *B. cereus* MSX-D12 *thsA* gene.

**Supplementary File 2**. Plasmid map of pGO2_thrC_Pxyl_dafuensis_ThsA, shuttle vector for *B. dafuensis* FJAT-25496 *thsA*

**Supplementary File 3**. Plasmid map of pGO3_amyE_hspank_cereus_ThsB, shuttle vector for *B. cereus* MSX-D12 *thsB* gene.

**Supplementary File 4**. Plasmid map of pGO4_amyE_hspank_dafuensis_TIR1, shuttle vector for *B. dafuensis* FJAT-25496 *tir1* gene.

**Supplementary File 5**. Plasmid map of pGO5_amyE_hspank_dafuensis_TIR2, shuttle vector for *B. dafuensis* FJAT-25496 *tir2* gene.

**Supplementary File 6**. Plasmid map of pGO6_amyE_hspank_GFP, shuttle vector for GFP control construct.

**Supplementary File 7**. Plasmid map of pGO7_amyE_hspank_dafuensis_TIR1+TIR2, shuttle vector for *B. dafuensis* FJAT-25496 *tir1+tir2* genes.

**Supplementary File 8**. Plasmid map of pGO8_pET30a_strep_BdTIR_His, protein expression plasmid for *Brachypodium distachyon* BdTIR gene.

**Supplementary File 9**. Plasmid map of pGO9_pAB151_cereus_ThsA_strep, protein expression plasmid for *B. cereus* ThsA purification.

### Supplementary figures

**Figure S1**. Effects of mutations in Thoeris genes on defense against phage SPO1.

**Figure S2**. NADase activity of ThsA following exposure to cell lysates and cADPR.

**Figure S3**. Production of cADPR isomer of by ThsB following phage infection.

**Figure S4**. Efficiency of plating on cells expressing Thoeris systems of *B. cereus, B. dafuensis* and hybrid systems.

### Supplementary Figures

**Figure S1.**
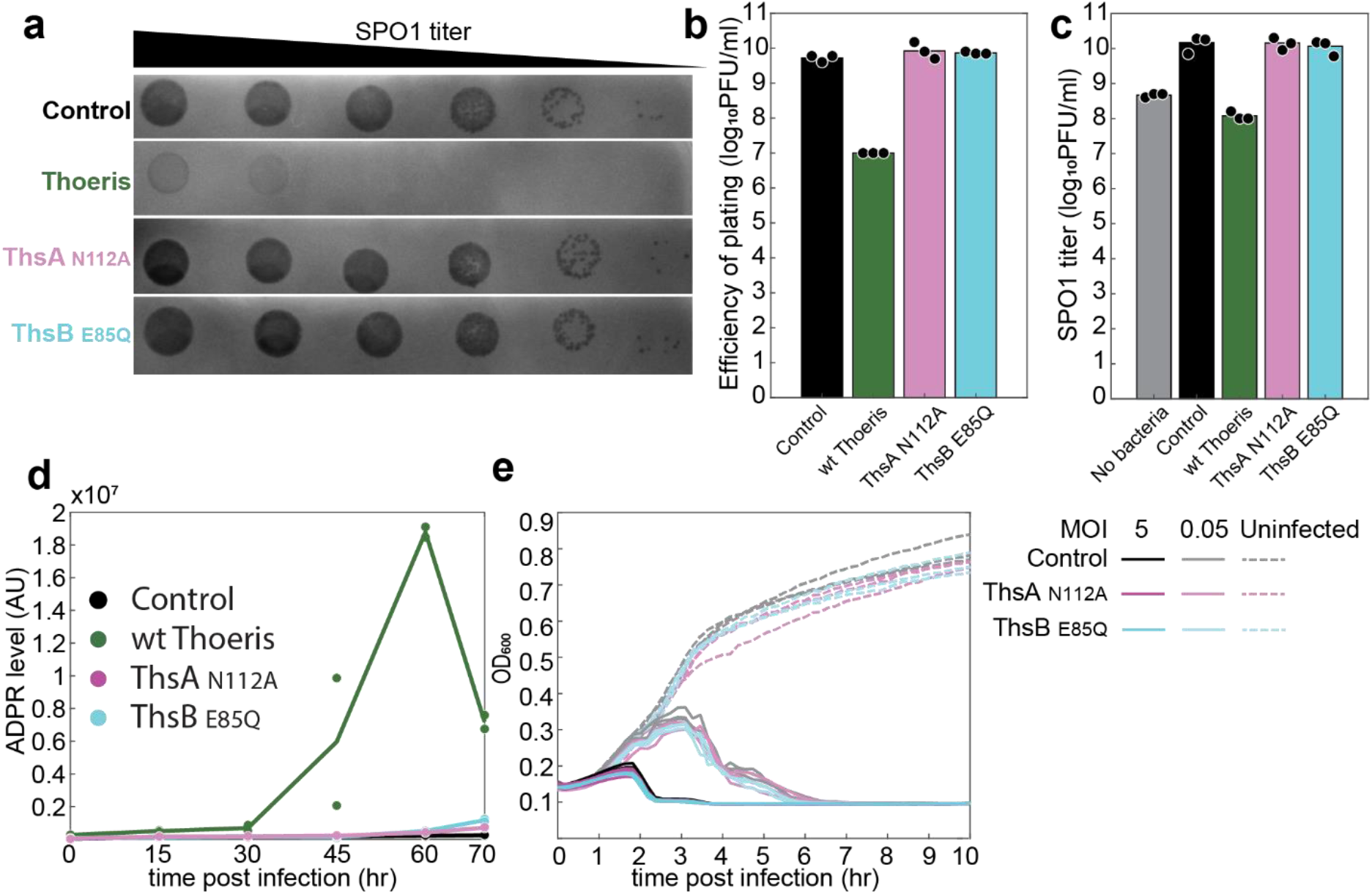
Effects of mutations in Thoeris genes on defense against phage SPO1. **(a)** Plaques of phage SPO1 on control cells (black), cells expressing both wt Thoeris proteins (green), cells expressing mutant ThsA_N112A_ and wt ThsB (magenta), or cells expressing wt ThsA and mutant ThsB_E85Q_ (cyan). Ten-fold serial dilution of the phage lysate were dropped on the plates. **(b)** Efficiency of plating (EOP) of phage SPO1 on control and Thoeris-containing strains, representing plaque-forming units per milliliter. Each bar graph represents average of three replicates, with individual data points overlaid. **(c)** Replication of phage SPO1 in the presence of Thoeris-containing and Thoeris-lacking (control) cells, or without cells (no bacteria). Lysates were collected 2.5 hours following infection of liquid cultures at an initial MOI of 5, and phage titer was quantified by plating serial dilution of the lysates on the control strain. **(d)** Adenosine diphosphate ribose (ADPR) levels in control culture (black), cells expressing wild type Thoeris (green), or mutated Thoeris systems (ThsA_N112A_ mutation in magenta and ThsB_E85Q_ in cyan) during infection. Time 0 represents uninfected cells. Cells were infected by phage SPO1 at an MOI of 5. Each line represents the mean of two independent experiments, with individual data points shown. ADPR levels were measured by LC-MS and calculated from the area under the curve of the identified ADPR peak at m/z 560.0805 (positive ionization mode) and retention time of 8.5 min. Peak was identified using a synthetic standard run. **(e)** Growth curves of Thoeris mutants during infection by phage SPO1 at MOI of 5 or 0.05. Curves represent three independent experiments.

**Figure S2.**
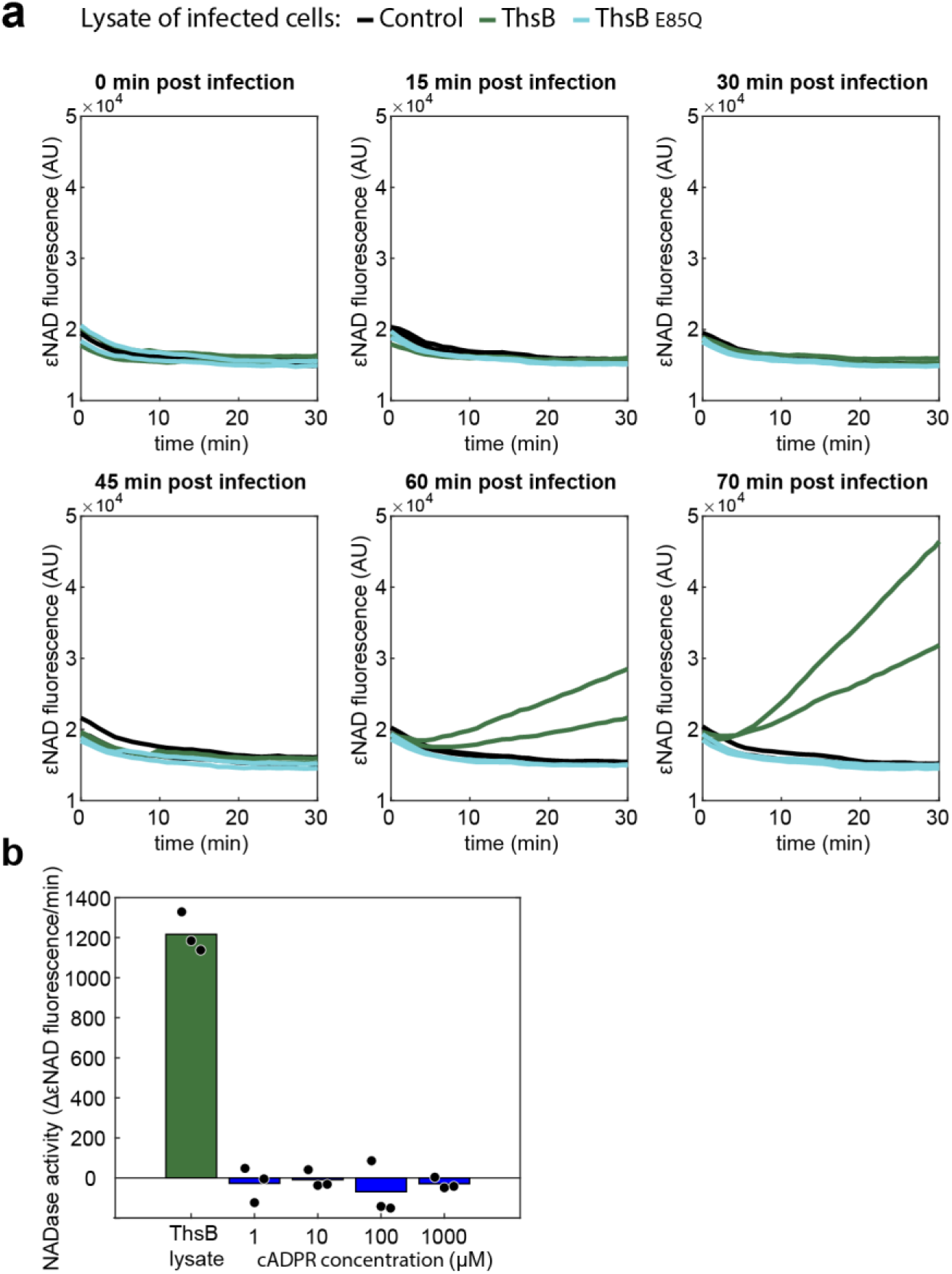
NADase activity of ThsA following exposure to cell lysates and cADPR. **(a)** εNAD fluorescence, indicating εNAD cleavage, of purified ThsA protein incubated with lysates derived from GFP-expressing control cells (black), ThsB-expressing cells (green) or cells expressing the mutant ThsB_E85Q_ (cyan), during infection. Each sub-panel represents lysates derived from a different time point following infection by phage SPO1 at MOI of 5. Time 0 represents uninfected samples. Data shown in Figure 2B represent rates calculated from the data shown here. **(b)** NADase activity of ThsA, when incubated with filtered cell lysates derived from ThsB-expressing cells 70-minutes post infection by phage SPO1, or with buffer containing 1 μM – 1000 μM of synthetic cyclic ADP-ribose (cADPR). NADase activity was calculated as the rate of change in εNAD fluorescence during the linear phase of the reaction. Bars represent mean of three experiments, with individual data points overlaid.

**Figure S3.**
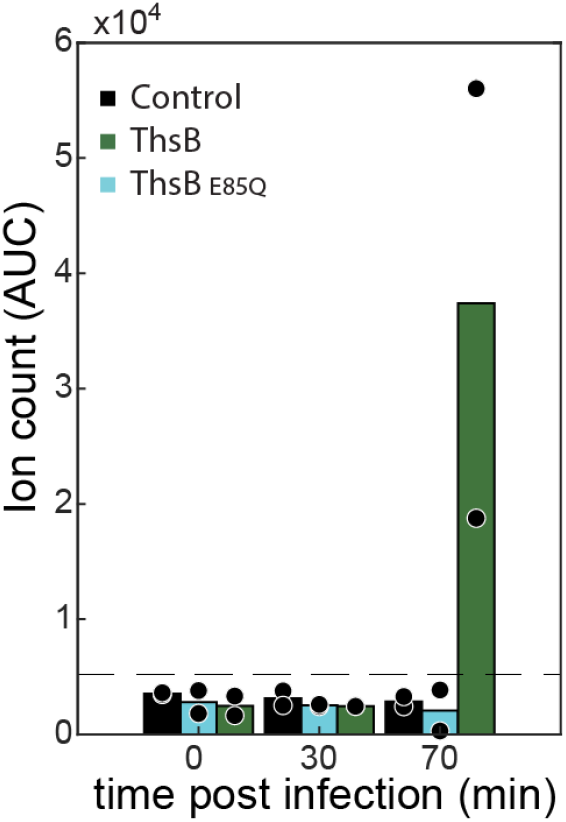
Production of cADPR isomer of by ThsB following phage infection. Detection of a cADPR isomer with an m/z of 542.0684 via LC-MS (positive ionization mode) in lysates derived from GFP-expressing (control), ThsB-expressing, and ThsB_E85Q_-expressing cells during SPO1 infection at MOI 5. Bars represent mean of two experiments, with individual data points overlaid. Dashed line represents limit of detection.

**Figure S4.**
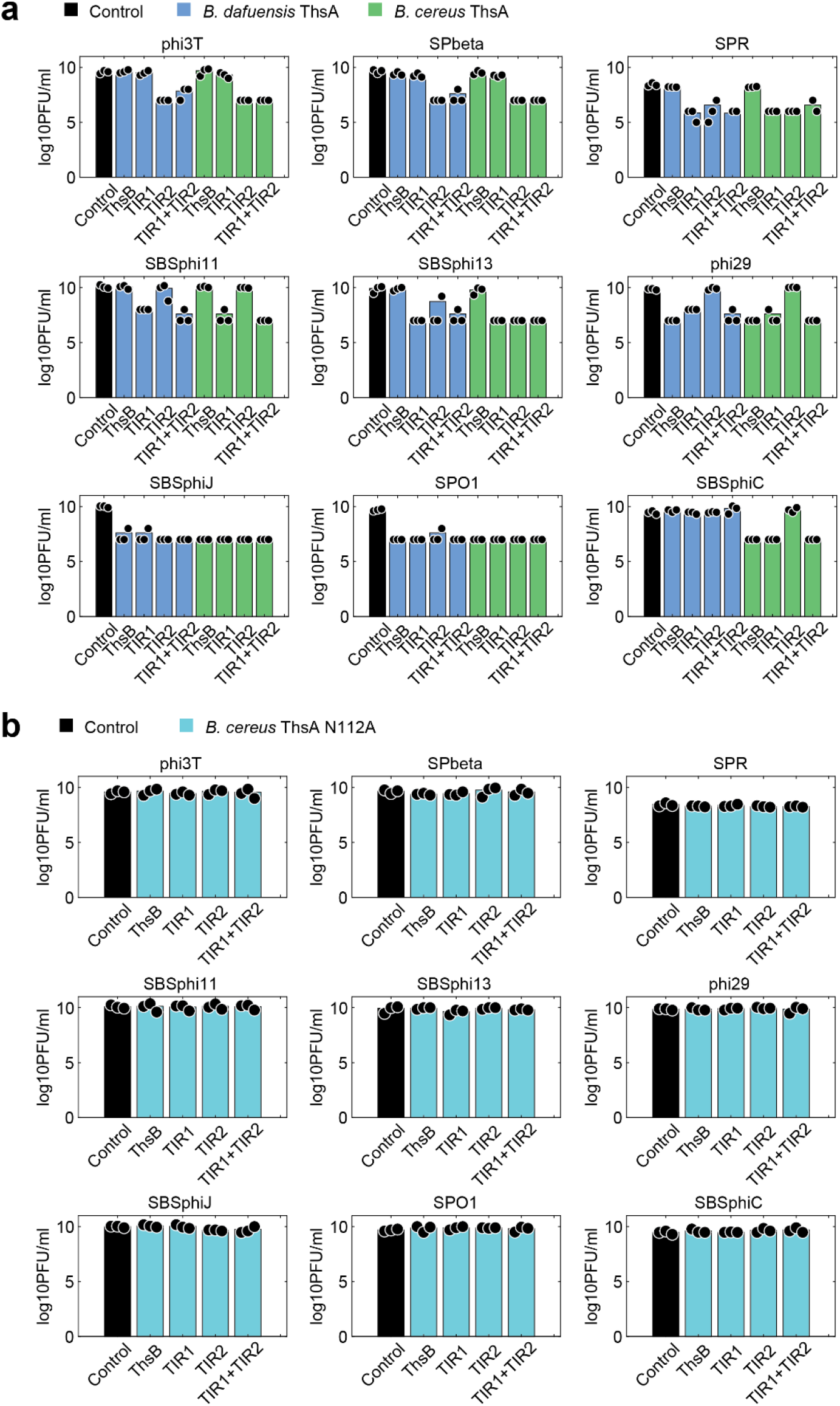
Efficiency of plating on cells expressing Thoeris systems of *B. cereus, B. dafuensis* and hybrid systems. **(a)** EOP of phages on control cells (black) and cells expressing combinations of ThsA from either *B. cereus* (green) or *B. dafuensis* (blue), TIR protein from *B. cereus* (ThsB), TIR proteins from *B. dafuensis* (TIR1 and TIR2) or both TIRs from *B. dafuensis* (TIR1+TIR2). Bars represent mean of 3 replicates, with individual data points overlaid. **(b)** EOP of phages on control cells (black) and cells expressing mutant ThsA_N112A_ from *B. cereus* together with combinations of TIR proteins from *B. cereus* and *B. dafuensis*.

